# Visualizing ATP Dynamics in Live Mice

**DOI:** 10.1101/2020.06.10.143560

**Authors:** Norimichi Koitabashi, Riki Ogasawara, Ryuto Yasui, Yuki Sugiura, Hinako Matsuda, Shigenori Nonaka, Takashi Izumi, Masahiko Kurabayashi, Makoto Suematsu, Motoko Yanagita, Hiromi Imamura, Masamichi Yamamoto

## Abstract

Analysis of the dynamics of adenosine triphosphate (ATP) is vital to quantitatively define the actual roles of ATP in biological activities. Here, we applied a genetically encoded Förster resonance energy transfer biosensor “GO-ATeam” and created a transgenic mouse model that allows systemic ATP levels to be quantitatively, sensitively, noninvasively, and spatiotemporally measured under physiological and pathological conditions. We used this model to readily conduct intravital imaging of ATP dynamics under three different conditions: during exercise, in all organs and cells; during myocardial infarction progression; and in response to the application of cardiotoxic drugs. These findings provide compelling evidence that the GO-ATeam mouse model is a powerful tool to investigate the multifarious functions of cellular ATP *in vivo* with unprecedented spatiotemporal resolution in real-time. This will inform predictions of molecular and morphological responses to perturbations of ATP levels, as well as the elucidation of physiological mechanisms that control ATP homeostasis.

**One Sentence Summary:** Intravital real-time imaging of ATP dynamics in multiple organs using GO-ATeam mice, can be used to quantitatively, sensitively, noninvasively, and spatiotemporally measure systemic ATP levels and provide a platform for preclinical pharmacological studies.

## INTRODUCTION

Multiple recent technologies, including RNA-Seq, have driven considerable advances toward a complete understanding and reliable prediction of biological activities at scales ranging from single cells to whole organisms (Wang et al., 2009). However, much is yet to be discovered regarding the coordinated and fluctuating biological activities in multicellular organs under physiological and pathological conditions, as they impact individual cells, because biological activities are regulated not only by intracellular signals but also by extracellular signals and the environment. Biological activities such as signal transduction, mRNA expression, and chromatin structure, as well as the activities of the proteins that regulate these processes, are all affected by intracellular adenosine triphosphate (ATP) levels (Fantl et al., 1993; Lusser and Kadonaga, 2003). ATP is also fundamentally important for many vital cellular processes such as maintaining the membrane potential and organelle transport in energy conversion (Dzeja et al., 2002) (Dzeja et al., 2003) (Kamerlin et al., 2013) (Magistretti and Allaman, 2015) (Zala et al., 2013). Thus, a quantitative analysis of *in vivo* ATP dynamics at the single-cell level can provide a means for investigating the dynamics of biological activities in multicellular tissues. Such an analysis could address the question of whether ATP levels may vary between different cell types within the same tissue, for example, or how much fluctuation is normal between individual cells of the same type. Historically, it was impossible to measure ATP levels of tissues with classical biochemical methods while preserving the integrity of organs with high spatiotemporal resolution (Khlyntseva, 2009). In recent years, methods for detecting ATP via UV−visible absorption (Jung et al., 2017), magnetic resonance spectroscopy (Befroy et al., 2012) (Chaumeil et al., 2009)or nuclear magnetic resonance (Guo et al., 2014) have been developed, but none of these can quantify ATP concentrations with high resolution at the single-cell level. In 2009 two genetically encoded fluorescent biosensors, called Perceval (Berg et al., 2009) and ATeam (Imamura et al., 2009), were invented and enabled imaging of the ATP/ADP ratio and ATP within living culture cells, respectively. Later, improved ATP biosensors and other types of ATP biosensors were reported, which include PercevalHR (Tantama et al., 2013), QUEEN (Yaginuma et al., 2014), MaLion (Arai et al., 2018), and GO-ATeam (Nakano et al., 2011). In this study, we chose a FRET-based ATP biosensor GO-ATeam, which employs green fluorescent protein as a donor and orange fluorescent protein as an acceptor, for studying ATP dynamics in living mammals. It is effective for these types of studies because of its minimal sensitivity to a broad pH range (6.3-8.3) and its ratiometric readout, which cancels fluctuation of fluorescent signals caused by movement of biological samples.

Here, we report the generation of an ATP visualization animal, “GO-ATeam mice”, in which the reporter achieve ubiquitous expression. Here we report our proof-of-principle analyses of different tissues and different methods. The ratio of FRET to GFP fluorescence intensities in intact cells was highly correlated with cytosolic ATP concentration determined by a proven biochemical method. These results prompted us to conduct imaging of live mice, for which we established GO-ATeam models of diverse physiological and pathological conditions. These findings support our expectation that GO-ATeam mice will serve as a useful platform for studying the dynamics of ATP *in vivo*, with the potential for conducting assays to elucidate the maintenance of energy homeostasis in physiology and possibly preclinical pharmacological studies.

## RESULTS

### Generation of Transgenic Mice to Determine ATP Dynamics *In Vivo*

We chose to employ the GO-ATeam strategy (Nakano et al., 2011) to observe ATP dynamics in live mice. As noted above, this system employs GFP and OFP as the FRET pair, which can be readily detected and is minimally sensitive to pH, an important advantage because metabolic stress can cause a drop in intracellular pH. After several failed attempts to generate GO-ATeam transgenic mice, we obtained stable GO-ATeam knock-in mice with the FRET reporter cassette (**Figs. 1A-1H)**. The knock-in mice yielded homozygous and heterozygous offspring consistent with Mendelian inheritance. Importantly, body weight, morphology and size, as well as the weights and functions of the organs, were normal. Moreover, the mice were phenotypically normal throughout their expected lifespan of approximately three years (data not shown).

**Fig. 1.**
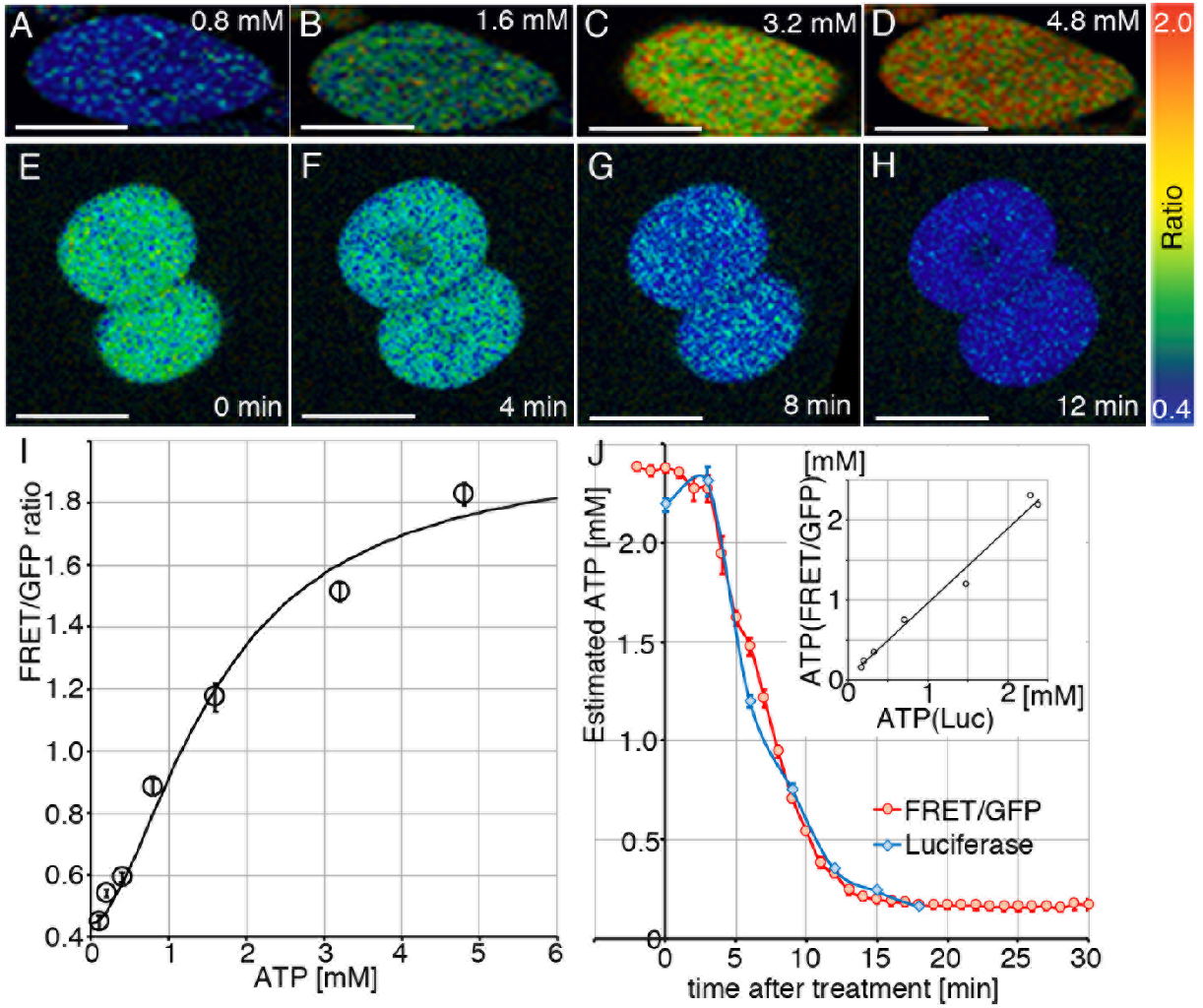
Measurement of Cytosolic ATP Levels in GO-ATeam transgenic mouse embryos and embryonic fibroblasts. (A-D) Images of FRET/GFP fluorescence emitted by a permeabilized mouse embryonic fibroblasts (MEF) derived from GO-ATeam2 knock-in mouse embryos incubated in calibration buffer. The ATP concentrations in the calibration buffer ranged from 0.8–4.8 mM. There was a close positive correlation between the FRET/GFP ratios and ATP concentrations from 0.1 mM to 6.0 mM (I) (n = 37). The plot was fitted with the Hill equation: 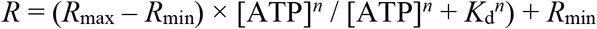, where *R*_max_ and *R*_min_ are the maximum and minimum fluorescence emission ratios, respectively, *K*_d_ is the apparent dissociation constant, and n is the Hill coefficient. (FRET/GFP) = (1.96 – 0.44) × [ATP]^1.7^ / ([ATP]^1.7^ + 1.6^1.7^) + 0.44. (E-H) FRET/GFP values calculated from images of two-cell embryos treated simultaneously with inhibitors of glycolysis and OXPHOS (2DG and antimycinA, respectively). The ATP concentrations estimated using the FRET/GFP ratio (n = 16) corresponded to those determined using the luciferase assay (J) (n = 118) (inset: R^2^ = 0.9846). Intensity-modulated display (IMD) images of the FRET/EGFP ratios (0.4 to 2.0) are shown. Scale bars indicate 50μm (A-D, I-J), or 25μm (E-H).

### The FRET/GFP Ratio Reliably Reflects Cytosolic ATP Concentrations in GO-ATeam Mice

To examine whether the GO-ATeam probe can effectively measure cytosolic ATP concentrations in GO-ATeam knock-in mice, we first investigated the fluorescence signals in mouse embryonic fibroblasts (MEFs) obtained from GO-ATeam knock-in mice. After permeabilization of the plasma membrane of MEFs (n = 37), we recorded FRET/GFP ratios in the cells with a two-photon microscope while stepwise increasing the ATP concentrations in the medium (**Figs. 1A-1D**). The intracellular FRET/GFP ratio changed as a function of applied ATP concentration, ranging from 0.1 mM to 6 mM. By fitting the dose-response plot with the Hill equation, we obtained a calibration curve for directly estimating ATP concentrations from the FRET/GFP ratios (**Fig. 1I**).

To further validate the FRET/GFP ratio as a quantitative measure of ATP concentration, we treated two-cell-stage embryos from knock-in mice with 2-deoxy-D-glucose to inhibit glycolysis, and antimycin A to inhibit OXPHOS, followed by estimations of ATP concentrations at certain time points either with fluorescence-based FRET imaging (**Figs. 1E-1H**, n = 16) or with a proven cell lysate-based firefly luciferase method (n = 118). The time-course of ATP change estimated by FRET imaging was virtually superimposable with the time-course obtained by the luciferase method (R^2^ = 0.9846) (**Fig. 1J**). Thus, we concluded that cytosolic ATP concentrations in living cells from GO-ATeam mice can be reliably estimated by imaging with a two-photon microscope.

We next examined whether the expression level of the GO-ATeam probe in the knock-in mice is sufficient for the quantitative measurement of FRET signals. To assess the sensitivity and accuracy of GO-ATeam FRET/GFP ratio measurements, we generated embryos expressing different levels of GO-ATeam2 by electroporating wild-type single-cell embryos with GO-ATeam2 mRNA (n = 756), and compared the estimated ATP concentrations from FRET/GFP ratios of the embryos acquired at 16 bits with a two-photon microscope (**Fig. S2,** red dots). The total average autofluorescence intensities of the embryos were 1389 ± 1.65 (n = 36). As expected, the estimated ATP concentrations of embryos showing low average fluorescence intensities varied widely. In contrast, those showing high average fluorescence intensities (>10,000, at least 7- fold higher than the autofluorescence intensities) were within a relatively narrow range (2.40 ± 0.01 mM), which were very close to the ATP concentration obtained from the luciferase assay (2.34 ± 0.06 mM, n = 49). The total average fluorescence intensities of all heterozygous and homozygous knock-in embryos were more than 10,000, and more than 7-fold higher than the autofluorescence intensities, indicating that the ATP level could be calculated accurately for these embryos. It should be added that the homozygous knock-in embryos had an average fluorescence intensity value of about twice that of the heterozygous knock-in embryos (n = 88 and n = 68, blue dots and green dots in Fig. S2, respectively). Therefore, as a rule, we accepted measurements with total average fluorescence intensities that were more than 7x the autofluorescence intensities in the analyses that follow.

In addition to a two-photon microscope, we also employed a fluorescence stereo microscope, which can capture low magnified features in the bodies. Fig. S1A-D shows the same images of liver slices from GO-ATeam mouse in normoxic and hypoxic conditions captured by a two-photon microscope and a fluorescence stereo microscope (**Figs. S1A–S1D**). The FRET/GFP ratios of the low magnified images were closely correlated with those of the two-photon micrographs (R^2^ = 0.92, n = 139, **Fig. S1E**), indicating that a fluorescence stereo microscope can be used for quantitative estimation of ATP levels in GO-ATeam mice at a low magnified scale.

### Differences in ATP Concentrations in Multiple Organs and Cell Types

In order to explore the heterogeneity in ATP concentrations within and between organs, we analyzed ATP levels in live neonatal (postnatal day 0) and adult (8 weeks of age) GO-ATeam mice. First, the animals underwent laparotomy after anesthesia, followed by imaging with a stereo microscope. The FRET/GFP images of the neonate (**Figs. 2A,B**) and those of the adult (**Fig. S3A**) revealed clear differences in ATP levels between organs. For example, brown adipose tissues, which were located between the shoulders, showed significantly lower FRET/GFP ratios compared with surrounding tissues (**Fig. 2A**). ATP concentrations of heart, lung, liver, kidney, pancreas, stomach, small intestine, and large intestine, which were estimated from fluorescence images, ranged approximately from 1 to 6 mM (**Fig.2C-J**, right column). We also estimated ATP concentrations of these organs using a firefly luciferase method (**Fig. 2C-J**, left column), and found that the values were roughly similar to those estimated from fluorescence images, except for the large intestine, although there was large variance in luciferase-based ATP concentrations. Because the penetration of the excitation light of the microscope into surface tissues steeply decreases, the stereo microscope only detects fluorescent signals from the surface of organs, while the luciferase method measures ATP in the whole organ. Thus, the large difference in estimated ATP concentrations of the large intestine observed between the two methods might suggest large variations in ATP concentrations within the organ; i.e., higher ATP in the muscle layer of the intestine, and lower ATP in the luminal tissues, such as villus. It is suggested that large variance is influenced by the time from organ harvest to luciferase assay and the efficiency of organ crushing.

**Fig. 2.**
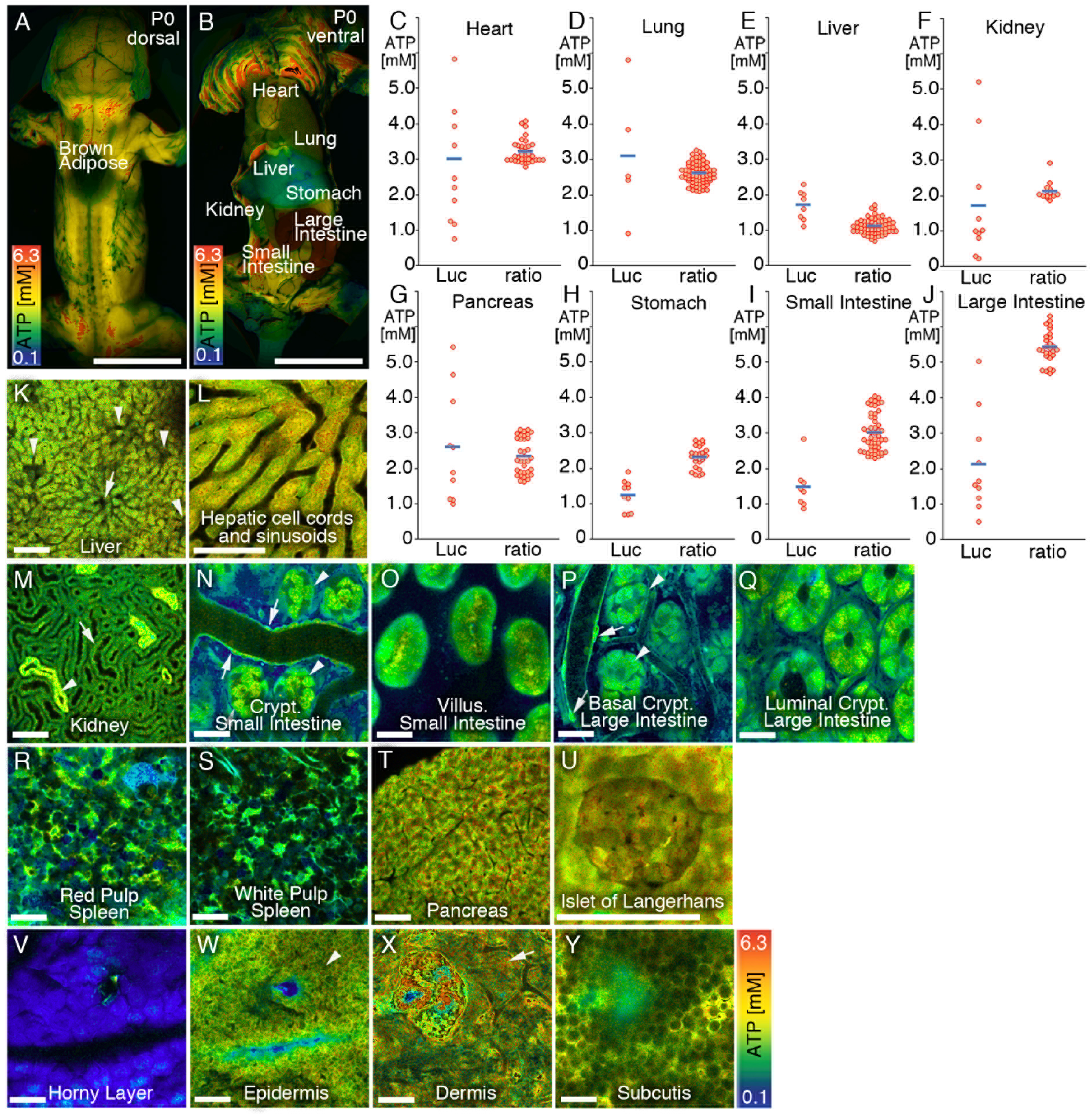
ATP levels in GO-ATeam Live Mice and Intravital Imaging of Organs. (A, B) FRET/GFP fluorescence ratios in postnatal day 0 ([A], dorsal; [B], ventral) GO-ATeam mice. (C–J) ATP concentrations measured using the luciferase assay (“Luc”) and FRET/GFP ratios (“ratio”) in heart (C), lung (D), liver (E), kidney (F), pancreas (G), stomach (H), small intestine (I), and large intestine (J) in neonatal GO-ATeam mice. (K–Y) Intravital FRET/GFP imaging of an adult (aged 8 weeks) GO-ATeam mouse. Liver (K, L, arrow, central venule; arrowhead, portal venule), kidney (M, arrow, proximal tubule; arrowhead, distal tubule), small intestine (N, O, arrow, blood vessel; arrowhead, paneth cells), large intestine (P, Q, arrow, blood vessel; arrowhead, paneth cells), spleen (R, S), pancreas (T, U), and skin (V–Y, arrow, epidermis; arrowhead, dermis). ATP concentrations (range, approximately 0.1 mM–6.3 mM) are depicted by the spectrum. Scale bars indicate 10mm (A, B), or 100μm (K-Y).

Next, we performed two-photon intravital FRET/GFP imaging to detect ATP in the deep tissues and cells of adult and neonatal GO-ATeam mice, which were anesthetized using intratracheal intubation. The analyses of adult (**Figs. 2K–2Y**) and neonatal (**Figs. S3J–S3O**) GO-ATeam mice included the following organs: liver (**Figs. 2K, 2L, and S3J**); kidney (**Figs. 2M, S3M, and S3N**); small intestine (**Figs. 2N, 2O, S3K, and S3L**); large intestine (**Figs. 2P and 2Q**); spleen (**Figs. 2R and 2S**); pancreas (**Figs. 2T, 2U, and S3O**; and skin (**Figs. 2V–2Y**). The images reveal striking differences in ATP concentrations among organs and cells.

### ATP Dynamics Associated with the Force Generated by Muscle Contraction

The role of ATP in fueling muscle contraction has been intensively studied for decades, mainly through investigations of small animals and only a few muscle types **(Barclay, 2017) (Barclay, 2015)**. However, it still unknown to what extent these previous studies on tissues, isolated muscle, or muscle cells—reflect the ATP dynamics in additional muscle types and other mammalian species with varying energy needs, because live animal imaging has not been exploited **(Barclay, 2015)**. *In vivo* analyses of ATP dynamics will likely enhance our understanding of the linkage between bioenergetics and muscle contraction. We began our *in vivo* approach by using GO-ATeam mice to study ATP dynamics associated with the force generated by muscle contraction.

For this purpose, we immobilized the legs of live mice and electrically stimulated the sciatic nerve to induce contractions of the tibialis anterior muscle (**Fig. S4A**). FRET/GFP ratios in the tibialis anterior muscle were recorded with the fluorescent stereo microscope (**Figs. 3A-J**), simultaneously with torques generated by the muscle contraction; a range of responses was generated by stimulating the sciatic nerve with various frequencies. The muscle underwent twitching constriction at 20 Hz stimulation, while tetanic constriction was observed at 100 Hz stimulation. After applying stimulation, immediate increases and decreases in torques (**Fig. 3K**) and FRET/GFP ratios (n = 4, **Fig. 3L**) were observed, respectively. Maximum torque was generated within 0.9 s upon 100 Hz stimulation (**Fig. 3K**). The usage of ATP expeditiously increased then decreased, even the peak torque was still being produced (**Fig. S4B**), implying that the production of force may become more efficient once muscle is maximally contracted. (Jones et al., 2009) reported that the ATP level present in all skeletal muscles before and after contractile motion is unchanged when examined by the firefly luciferase method. However, when the identical type of anterior cervical muscle was measured before and after contraction using the ATP visualization mouse technique, the ATP level decreased with contraction (**Fig. 3**). The effect was subtle, as the ATP level decreased by only 0.6 mM even at 100 Hz exercise. These results suggest that the firefly luciferase method did not detect the decrease in ATP levels due to limitations of the experimental technique. The changes in both the torques and FRET/GFP ratio were highly dependent on the frequency of the applied stimulations. The simultaneous measurement of ATP levels and torque in real time showed that considerable amounts of ATP were consumed when generating torque, and that while maintaining torque the same amount of ATP was used regardless of the magnitude of the torque. These results demonstrate the utility of the GO-ATeam mouse for quantitative evaluations of energy efficiency in muscle strength (i.e. torque) of various muscles; in addition, it will help to clarify the differences between sarcomeres that occur unevenly in muscle cells, and which might vary with the distribution and timing of energy use.

**Fig. 3.**
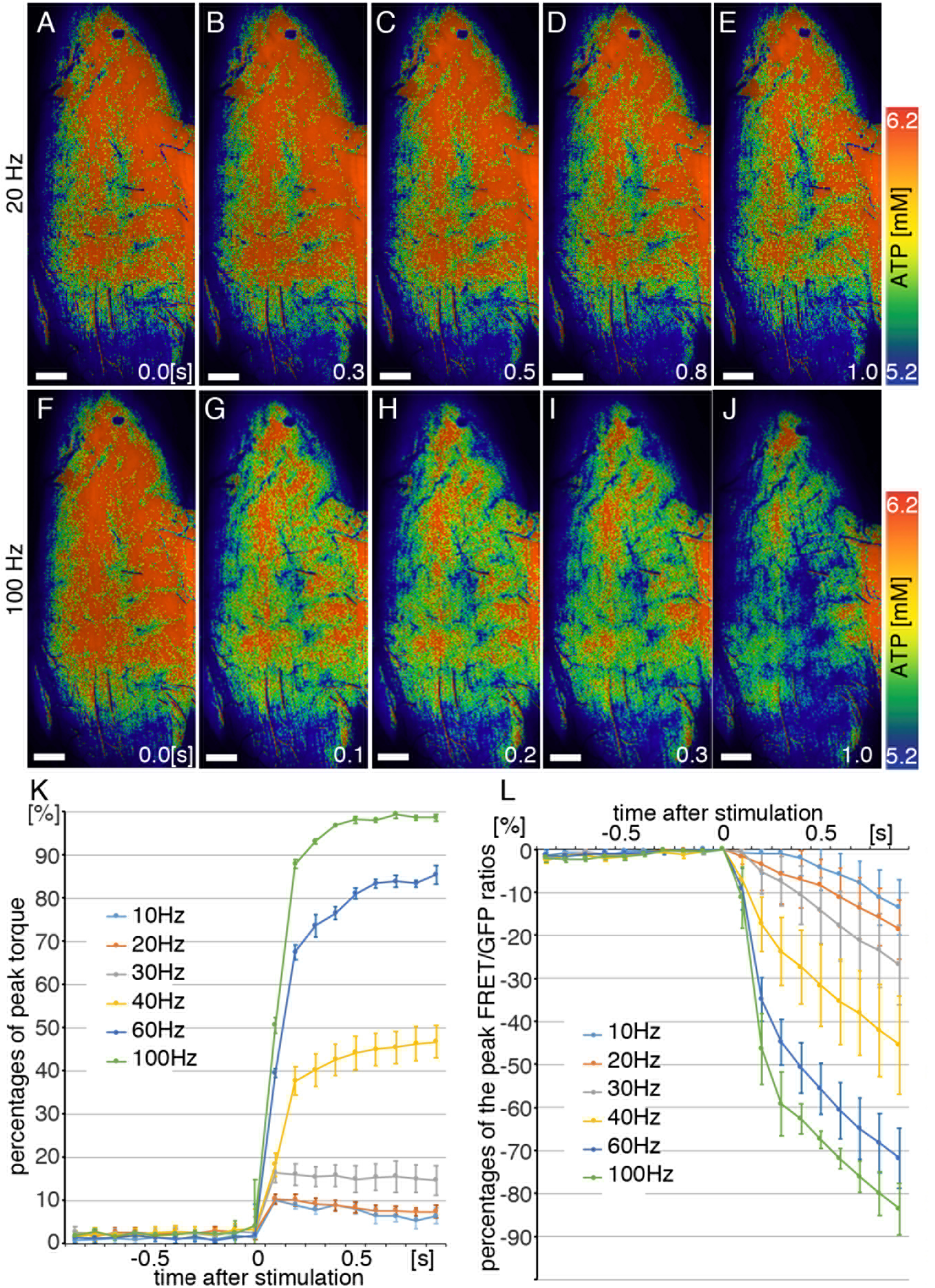
ATP Dynamics Correspond to the Force Generated by Muscles. Intravital imaging of FRET/GFP ratios of the tibialis anterior muscle subjected to stimulation of the sciatic nerve (A–J). Stimulation frequencies ranged from 20 Hz (A–E) to 100 Hz (F–J). The percentages of peak torque (K) (n = 4) and the percentages of the peak FRET/GFP ratios (L) (n = 4) as a function of time after stimulating the sciatic nerve. The differences between frequency intervals differed significantly (L), except for 10 Hz and 20 Hz (K). Numbers indicate seconds after the stimulation, scale bars indicate 100μm (A-J).

### The GO-ATeam Reporter Facilitates Studies of the Local and Peripheral Effects of an Acute Pathological Insult

We next sought to determine whether GO-ATeam mice serve as an accurate and sensitive reporter of the local and global effects on ATP dynamics of an acute pathological insult of major medical significance. We were particularly interested in our ability to monitor ATP dynamics in organs and tissues peripheral to the primary site of pathology, because such information may aid in diagnosis at early stages of disease that are otherwise difficult to detect. In humans, the heart is the organ that consumes the most ATP, and ischemic heart disease is the leading cause of death worldwide (Opie, 2003). We created a myocardial infarction model by ligating the left anterior descending artery (LAD) in GO-ATeam mice. We acquired intravital images of ATP dynamics in sham-operated (n = 8) (**Figs. 4A, 4C, 4E, S5A, and S5C**) and experimental mice (n = 8) (**Figs. 4B, 4D, 4F, S5B, and S5D**) 5 days after the procedure. At this time, echocardiography revealed marked left ventricular dysfunction with focal hypokinesis and enlargement of the heart chamber, and histology revealed the infarction scar (data not shown) at the anterior wall. Intravital FRET/GFP imaging showed that ATP concentration was clearly diminished in the ischemic region of the LV, as well as in other organs such as the liver, large intestine, kidneys, and small intestine (**Figs. 4B, 4D, 4F, S5B, D**). On the other hand, ATP levels increased around the ischemic region of the LV. This increase in ATP levels within the “border zone” is consistent with the fact that energy charge rises around the infarct lesion which was previously shown in heart and brain ischemia models using imaging mass spectrometry(Hattori et al., 2010) (Sugiura et al., 2016). These results indicate that the use of GO-ATeam mice can be used to analyze the ATP dynamics throughout the body on a time scale up to several days.

**Fig. 4.**
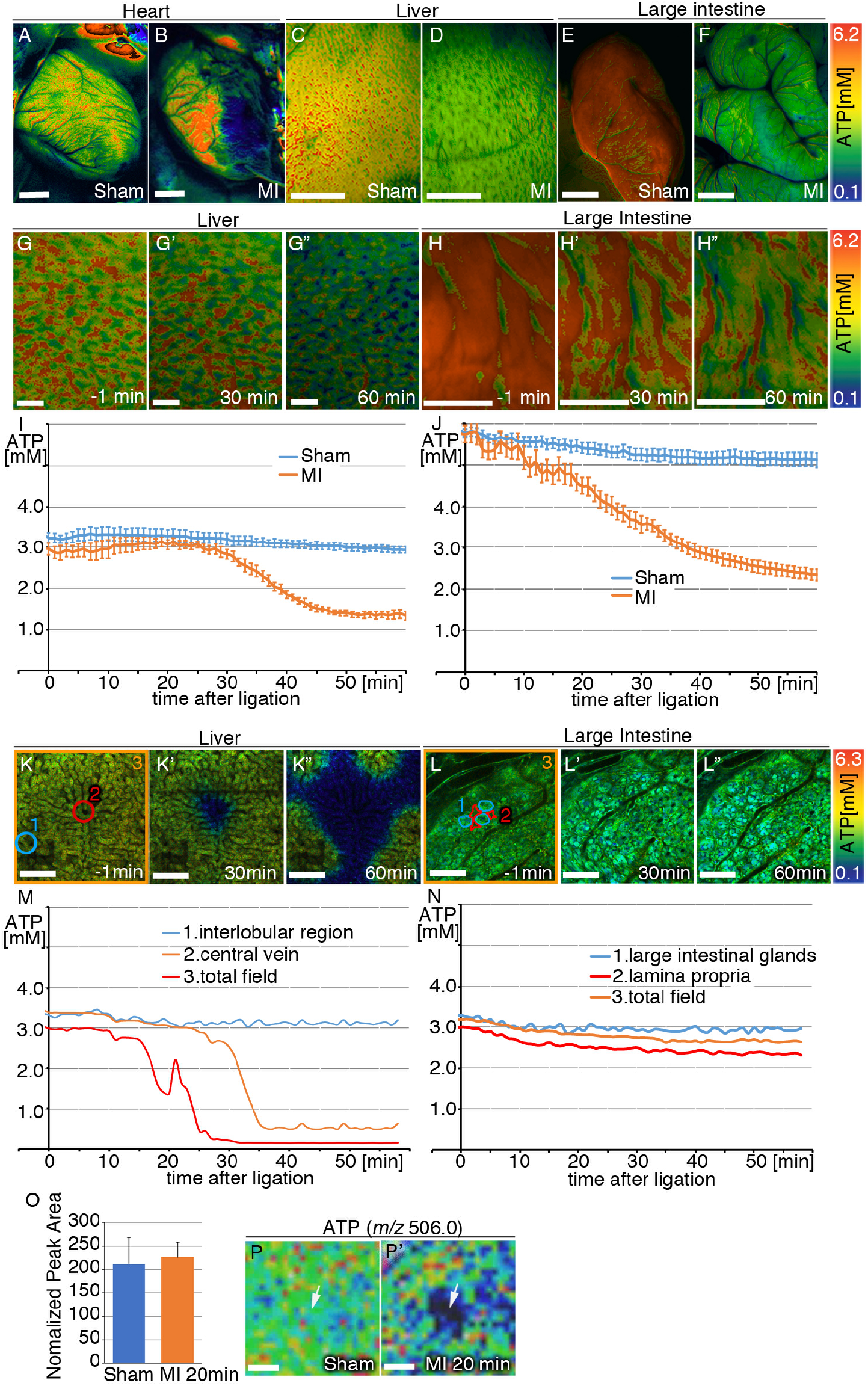
Effects of Myocardial Infarction on ATP Concentrations in the Organs of GO-ATeam Mice. (A–F) Intravital low magnified imaging of FRET/GFP ratios 5 days after ligation of the left anterior descending artery (LAD) (myocardial infarction, MI) (B, D, F; n = 8 each) or sham-operated mice (sham) (A, C, E; n = 8 each). Intravital time-lapse imaging of ATP concentrations calculated from the FRET/GFP ratios in the liver and large intestine using the fluorescence stereo microscope (G–G”, H–H”; I [graph of G-G”]; J [graph of H-H”], sham-operated, and MI; n = 9 and n = 6, n = 9 and n = 6, respectively, after t = 31 min; p<0.05, after t = 13 min, p<0.05) and respective cells using a two-photon microscope (K–K”, L–L”; M [graph of K-K”]; N [graph of L-L”]) after ligation of the LAD. (O-P’) Imaging Mass Spectrometry of ATP (*m/z* 506.0) in a sham-operated liver and a liver 20 min after ligation of the LAD (O, P, P’: arrow, central vein). In Figs. K and M, the regions of interest (ROIs) 1 to 3 show the periphery of the interlobular region, central vein, and total field of view, respectively. In Figs. L and N, ROIs 1 to 3 show the large intestinal glands, lamina propria, and total field of view, respectively. Scale bars indicate 2mm (A-F), 1mm (G-H”), or 100μm (K-L”, P, P’).

We next used the stereo microscope to perform time-lapse imaging of the whole body immediately after the LAD ligation to examine minute-scale changes in the whole body during acute heart failure induced by myocardial infarction. We found that the ATP levels in the liver (n = 6) (**Figs. 4G–4G” and 4I**) and kidney (n = 4) (**Figs. S5E– 5E”**) started to decrease (P < 0.05) after 31 min and 32 min after LAD ligation, respectively. ATP levels in the large intestine (n = 6) (**Figs. 4H–4H” and 4J**) and small intestine (n = 6) (**Figs. S5F–5F” and S5I**) started to decrease (P < 0.05) much earlier, 13 min and 12 min minutes after LAD ligation, respectively. Blood flow in the liver decreased by approximately 60% after ligation of the LAD (data not shown). These results indicate that GO-ATeam mice can be used to analyze global effects on ATP dynamics in response to acute pathological insults.

Next, we used two-photon microscopy to investigate the LAD ligation-induced alterations in ATP dynamics of organs at the cellular level. We found that ATP levels after 10 min only decreased around the pericentral regions in the liver (**Figs. 4K–K” and 4M**; ROI 2, red), while ATP levels were maintained along the periphery of the interlobular region including periportal regions (**Figs. 4K–K” and 4M**; ROI 1, blue), showing intralobular heterogeneity in ATP drop in response to hypovolemic hypoxia. This observation is consistent with the previous report showing greater susceptibility of pericentral regions to hypoxia (Suematsu et al., 1992a; Suematsu et al., 1992b). In the large intestine, ATP levels gradually decreased in the large intestinal glands and lamina propria (**Figs. 4L–L” and 4N**; ROIs 1, 2 and 3, blue, red and orange). Glands tended to maintain ATP levels compared with the lamina propria. These results demonstrate that there is a large intercellular heterogeneity in the reduction of ATP during hypoperfusion and hypoxia, even within the same organ, in real-time analysis. Clinically, for example, hepatocyte necrosis is observed near the central vein on a time scale of days after myocardial infarction. Therefore, although the time scales are different, these data show the order and area of ATP reduction in each of these organs correlates with clinical information on organ abnormalities and necrosis during myocardial infarction (Sherlock, 1951).

We also detected a decrease in the amount of ATP within a limited region near a central vein in the liver by mass spectrometry imaging 20 min after LAD ligation (**Figs. 4P**), consistent with the above FRET observation. In contrast, metabolome analysis of a whole liver did not detect the ATP decrease at the same time point (Fig. 4O). Thus, ATP imaging using GO-ATeam mice can detect local bioenergetic changes in real time with high sensitivity.

### Assessment of Drug-Induced Cardiotoxicity in GO-ATeam2 Mice

Drug-induced cardiotoxicity is a significant safety issue in drug development because it can be fatal (Watkins, 2011). However, cardiotoxicity may become apparent after clinical trials and marketing, and risk assessment in nonclinical trials has been difficult.

The mouse heart contracts and relaxes about seven times per second using a large amount of ATP. This exquisite balance between supply and consumption keeps the amount of ATP in the cytoplasm of cardiomyocyte constant. However, if the balance is slightly disturbed due to the toxicity of a drug to the heart, ATP concentrations will change within a short time. Thus, we hypothesized that cardiotoxicity of a drug may be detected as a change in the ATP dynamics of the heart before a change appears on the electrocardiogram, etc.(**Fig. 5A**). To test this, we examined anticancer drugs with reported cardiotoxicity, antiarrhythmic drugs and antibiotics that induce torsade point (TdP).

**Fig. 5.**
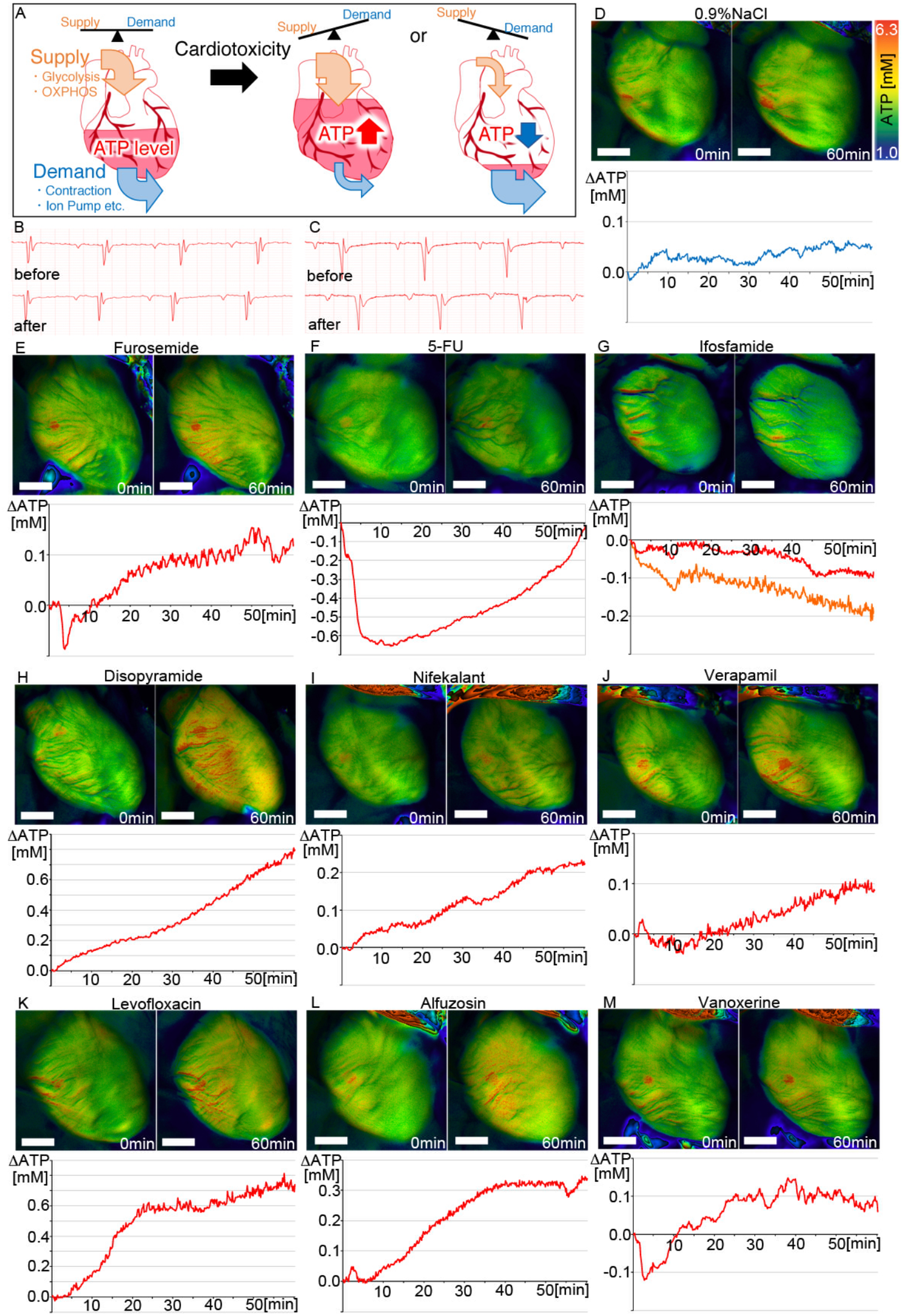
Effects of Drug-Induced Cardiotoxicity on ATP Concentration in the Heart of GO-ATeam Mice. (A) Scheme: Cardiac ATP levels are balanced by supply (glycolysis, OXPHOS, etc.) and demand (contraction, ion pumps, etc.). Due to the drug, cardiotoxicity causes an imbalance between ATP supply and demand, and is expected to alter cardiac ATP levels in a short time. (B, C) electro-cardiogram. The identifier “before” refers to before administration, and “after” refers to 1 hour after administration of the drug, 0.9% NaCl (B) or disopyramide (C). (D-M) Intravital time-lapse imaging of ATP concentrations calculated from the FRET/GFP ratios in the heart using the fluorescence stereo microscope (upper left, before administration; upper right, 60 minutes after administration, with graphical representations shown below each set of panels). The graphs show the change volume in ATP level of whole heart (y-axis) after administration (blue line, 0.9% NaCl; red line, indicated drug; orange line, ifosfamide in left ventricle). Horizontal axis shows time [minutes] after administration. (D) 0.9% NaCl (n=10). (E) furosemide (n=9). (F) 5-FU (n=6). (G) ifosfamide (n=5). (H) disopyramide (n=6). (I) nifekalant (n=6). (J) verapamil (n=6). (K) levofloxacin (n=5). (L) alfuzosin (n=5). (M) vanoxerine (n=4). Scale bars indicate 2mm (D-M).

Cardiac ATP levels were observed for 1 hour using a fluorescence stereo microscope while the drug was continuously administered via the jugular vein at concentrations that did not elicit an abnormal electrocardiogram (**Figs. 5B, C**). Examination of the time-course changes in the ATP level in the heart immediately before administration showed that there was almost no change in physiological saline (n= 10) and furosemide (n=9), a diuretic with no cardiotoxicity, with a rise of about 0.05-0.1 mM, consistent with the data shown in **Fig. 1** (**Figs. 5D and E**). In contrast, administration of doxorubicin (an anthracycline anticancer drug, n=3), 5-FU (an antimetabolite anticancer drug, n=6), and cyclophosphamide (an alkylating agent, n=4), rapidly reduced ATP levels, which then recovered (**Figs. 5F, S6A and S6B**). This is consistent with reports that doxorubicin accumulates in mitochondria in cardiomyocytes, causing increased oxidative stress and mitochondrial dysfunction (Ichikawa et al., 2014; Zhang et al., 2012). It is also consistent with clinical reports that 5-FU causes transient coronary vasospasm and cardiac ischemia, and that cyclophosphamide causes myocardial damage (Schimmel et al., 2004). On the other hand, when the alkylating agent ifosfamide was administered, the ATP level decreased only moderately in the entire heart, but the ATP level decreased significantly only in the left ventricle (**Fig. 5G**, n=5). This is consistent with clinical reports of ifosfamide eliciting left ventricular dysfunction (Cardinale et al., 2000). These results suggest that the cardiotoxicity of each anticancer drug can be detected as a change in ATP dynamics, adding consistent molecular evidence to clinical reports.

In addition to anticancer drugs, we evaluated the drug-induced TdP, a fatal ventricular arrhythmia as a measure of cardiotoxicity. TdP has been a significant safety issue in drug development; current predictions of TdP in non-clinical trials are not fully consistent with clinical TdP inducibility (Laverty et al., 2011).

We hypothesized that energy abnormalities might be related to myocardial electrical instability. Therefore, GO-ATeam2 mice were used to determine whether the results of the comprehensive *in vitro* proarrhythmia assay CiPA (Strauss et al., 2019) could be improved. First, the antiarrhythmic drugs disopyramide, procainamide, nifekalant, verapamil, and vanoxerine were examined (**Figs. 5H-J, 5M and S6C**). Disopyramide has been reported to reduce myocardial contractility by blocking Na^+^ channels (Mathur, 1972). The administration of TdP-inducing antiarrhythmic drugs disopyramide, procainamide and nifekalant all increased the ATP level in the heart by 0.25 mM or more by continuous administration for 60 minutes (**Figs. 5H, I, S6C**, n=6, 6, 5). On the other hand, verapamil, an antiarrhythmic drug that blocks hERG channels but suppresses TdP, showed little change in ATP levels (**Fig. 5J**, n=6) (Milberg et al., 2005 124). Similarly, vanoxerine, which blocks hERG channels but does not affect QT prolongation in the heart, did not alter ATP levels (**Fig. 5M**, n=4) (Lacerda et al., 2010). Next, antibiotics and antifungals such as levofloxacin, erythromycin, amphotericin B, azithromycin, ciprofloxacin, and metronidazole were examined (**Figs. 5K, S6E-I**, n=5, 7, 4, 5, 5, 3). As with antiarrhythmic drugs, continuous administration for 60 minutes increased the intracardiac ATP level by 0.25 mM or more for all antibiotics and antifungals, despite having different actions and chemical structures (**Figs. 5K, S6E-I**, data not shown). Furthermore, continuous administration of alfuzosin (n=5), a prodrug for inducing TdP, and droperidol (n=9), an antipsychotic, increased intracardiac ATP levels by 0.25 mM or more by continuous administration for 60 minutes (**Figs. 5L, S6**). Of the approximately 60 TdP-inducing drugs registered with the FDA, all 11 that we tested raised ATP levels by 0.25 mM or more by continuous administration for 60 minutes. These results indicate that the GO-ATeam mouse model can reliably identify drugs that induce TdP based on changes in the amount of cellular ATP.

## DISCUSSION

### Adaptable to Other Imaging Technologies

In the GO-ATeam2 mouse model, identification of cell types and observation of cell morphology can be observed by expressing far-red fluorescent proteins with different spectra, such as mCardinal (Chu et al., 2014) and mRaspberry (Wang et al., 2004), in the nucleus and cell membrane. A chemical dye can provide the fluorescent signal for a third color. Specific cell labeling using DiD and ER labeling using ER-Tracker Blue-White DPX can also be performed. ATP levels can also can be monitored simultaneously with the fluctuation of mitochondrial mass using MitoTracker DeepRed. Since the GO-ATeam mouse model can be used for a simple allele knock-in of GO-ATeam2, it can be similarly modified by crossing with other transgenic mice and various genetically modified mice. Examples include the GO-ATeam Amyotrophic lateral sclerosis model and the heart failure model.

In the GO-ATeam mouse model, spatiotemporal information on ATP dynamics is obtained from the organ level to the cell level in the whole mouse within the same individual after the onset of the disease (myocardial infarction) or after drug administration. It can be applied not only to intravital imaging but also to ATP dynamic observation under various conditions such as *ex vivo* and primary cultured cells, such as organ slices.

### Indicator of ATP Levels

Cellular ATP levels and ATP sensing are integral to an assortment of regulatory processes, including phosphorylation of signaling molecules, epigenetic factors, chromatin remodeling factors, and activation of ion pumps (Lusser and Kadonaga, 2003) (Fantl et al., 1993) (Skou, 1965) (Becker and Horz, 2002). Feedback mechanisms can influence enzymatic activity in response to changes in ATP concentration. Thus, fluctuations in ATP levels are can indicate a change in an organ or cell function that is not simply correlated with the amount of these proteins. Turning this around, knowing how ATP concentrations affect biochemical enzymatic activities may allow the direct measurement of ATP levels to serve as a proxy for biochemical assays. This implies that if ATP dynamics are quantified in real time, *in vivo*, spatiotemporal information related to functional changes can be obtained at the cellular level. For example, in cortical neurons, ATP levels are involved in the depth of the resting membrane potential and control nerve firing, because ATP is required for ion pumps, and the ATP concentration correlates with ion pump activity (manuscript in preparation). It will be interesting to compare ATP levels with gene expression profiles obtained from comprehensive analyses such as RNA-Seq at the cell level, and correlate these with spatiotemporal information. We presume that the main cellular factors controlled by the ATP level differ depending on the organ in question, cell type, and the environment. However, it is necessary for the near future to clarify these major factors by artificially increasing or decreasing ATP levels at the cellular level *in vivo*.

### A model for evaluating drug effects on ATP homeostasis in living animal

In general, ATP levels in the cytoplasm are always kept constant by balancing between consumption and supply in living cells (Ingwall, 2004). On the other hand, if there is a spatiotemporal perturbation of the consumption/supply balance, the ATP levels in the cytoplasm are expected to change. Glycolysis and OXPHOS are responsible for the generation of ATP. However, OXPHOS is much more efficient at ATP production than glycolysis, so most ATP production in normal cells depends on OXPHOS. If the ATP supply from OXPHOS decreases due to functional decline, such as mitochondrial injury, the supply is temporarily compensated for by using ATP reserves inside the cell, and ATP consumption is reduced. Subsequently, the body maintains homeostasis by activating the glycolysis system to restore the total supply of ATP and rebalance the energy homeostasis. (Ingwall, 2009). In the initial stage of heart disease, either metabolic stress (e.g., ischemia) or mechanical overload (e.g., pressure-overload) alters the ATP homeostasis (Kolwicz et al., 2013). Long-term metabolic stress to maintain an appropriate intracellular ATP level as above leads to cellular dysfunction, cell death, and heart failure (Kolwicz et al., 2013). To understand this progression, monitoring the spatiotemporal changes in ATP levels would be informative. In addition, for anti-cancer drug-induced cardiomyopathy, ATP homeostasis and mitochondrial function are key as well (Wallace et al., 2020). As a proof of principle, ATP visualization experiments with drug administration were performed here. All the drugs tested to induce cardiotoxicity also changed ATP levels in the heart in a short time, as observed with GO-ATeam FRET (**Fig. 5, S6**). We speculate that anti-cancer drugs appear to decrease cardiac ATP production due to mitochondrial damage. The mitochondrial damage and decreased OXPHOS may increase myocardial oxidative stress and irreversible damage in late phases of anti-cancer treatment (Wallace et al., 2020). ATP changes elicited by antiarrhythmic drugs are qualitatively as well as quantitatively different from those produced by anticancer drugs. Antiarrhythmic drugs cause “negative inotropic effects” of contractile proteins, thereby reducing ATP consumption. Since ATP imaging reflects the difference between ATP production and consumption, ATP imaging in the beating heart showed that the intracellular ATP level increases in response to antiarrhythmic drug administration. Conceivably, ATP production from mitochondria might be increased by anti-arrhythmic drugs, although the hypothetical mechanism is unknown. In any case, the increase in the intracellular ATP level may lower the membrane potential and thereby cause an arrhythmogenic effect.

Our data show that GO-ATeam mice can elucidate physiological phenomena by accurately measuring ATP levels over time in the cells and tissues *in vivo*. The GO-ATeam system can potentially be extended to a wide range of applications, such as elucidating networks between organs throughout the body and assessing toxicity.

## ACKNOWLEDGMENTS

We thank Mai Samejima and Takashi Musashi for their assistance in cardiac ATP imaging analysis, the members of Motoko Yanagita’s laboratory for assistance with experiments and useful discussions, Yoshitaka Ishihara for electroporation, Eiichiro Uchino for assistance with data analysis.

## Funding

This work was supported by Grant-in-Aids for Scientific Research (KAKENHI Grant Number 16K09491 to N.K.), the Japan Agency for Medical Research and Development (AMED Grant Number 19gm1210009, JP19gm5010002 and JP19gm0610011); partially by grants from KAKENHI (26293202, 17H04187), Grant in Aid for Scientific Research on Innovative Areas “Stem Cell Agingand Disease (17H05642)”, “Lipoquality (18H04673)” (all fonding to M. Yanagita), the JST, PRESTO (JPMJPR14MF), JSPS (KAKENHI Grant Number JP24116703), AMED (Grant Number JP17gm5010002), the Naito Foundation, the Mitsui Life Social Welfare Foundation, the Takeda Science Foundation, the Mother and Child Health Foundation, the Uehara Memorial Foundation, the Japan Brain Foundation, the Nakajima Foundation, the Foundation for Dietary Scientific Research, and the Lotte Foundation (all funding to M.Yamamoto).

## Author contributions

N.K., R.O. and M. Yamamoto designed the experiments. M. Yamamoto supervised the project. N.K., R.O., R.Y., Y.S., H.A. and M. Yamamoto performed experiments. N.K., R.O., R.Y., Y.S., H.A., S.N. and M. Yamamoto analyzed the data. All authors discussed the results. H. I. and M. Yamamoto wrote the manuscript.

## Competing interests

M.Yamamoto is involved in a pending patent related to GO-ATeam mice. H.I. holds a patent for ATeam probe. All other authors declare no competing interests.

## Data and materials availability

All data is available in the main text or the supplementary materials.

## LIST OF SUPPLEMENTARY MATERIALS

KEY RESOURCES TABLE

CONTACT FOR REAGENT AND RESOURCE SHARING

EXPERIMENTAL MODELS AND SUBJECT DETAILS

o Imaging of Muscle Contraction and Measurement of Torque
o GO-ATeam Mouse Model of Myocardial Infarction
o Drug-Induced Cardiotoxicity in GO-ATeam Mice

METHOD DETAILS

o Cell/Tissue Culture
o Embryo Culture
o Live Imaging of Embryos
o Intravital Imaging of Organs
o Metabolome analysis
o MS imaging
o Organ slices
o Image Processing
o Measurement of ATP Levels in Mouse Embryonic Fibroblasts (MEFs)
o Luciferase Assay
o Electroporation

QUANTIFICATION AND STATISTICAL ANALYSIS

DATA AND SOFTWARE AVAILABILITY

SUPPLEMENTAL FIGURE LEGENDS

