## Supplementary materials and figures for "Visualizing ATP Dynamics in Live Mice"

### KEY RESOURCES TABLE

| **REAGENT OR RESOURCE** | **SOURCE** | **IDENTIFIER** |
|---|---|---|
| **Chemicals, Peptides, Recombinant Proteins, Reagents, Kits** | | |
| 2-deoxy-D-glucose | **Wako** | **Cat#040-06483** |
| 5-FU | **Kyowa Kirin** | **Cat#46629324** |
| alfuzosin hydrochloride | **Tocris** | **Cat#3305/10** |
| alpha-hemolysin | **Sigma** | **Cat#H9395** |
| amphotericin B | **Sumitomo Dainippon** | |
| antimycin A | **Sigma** | **Cat#A8674** |
| Aron Alpha A | **Daiichi-Sankyo** | **Cat#13000494** |
| AsiSI | **NEB** | **Cat#R0630** |
| azithromycin | **Pfizer** | **Cat#46468675** |
| ciprofloxacin | **Meiji** | |
| cyclophosphamide | **Shionogi** | **Cat#11016024** |
| disopyramide | **Sanofi** | |

| | | |
|---|---|---|
| DMEM | **ThermoFisher** | **Cat#12100046** |
| DMSO | **Nacalai tesque** | |
| doxorubicin | **Nippon Kayaku** | |
| droperidol | **Daiichi-Sankyo** | |
| erythromycin | **Mylan** | |
| FBS | **JRScientific** | **Cat#43635** |
| furosemide | **Nichi-Iko** | |
| gDNA Eraser | **Takara** | **Cat#RR047A** |
| GeneArt Seamless Recombination System | **Thermo Fisher** | **Cat#A13288** |
| HEPES | **Sigma** | **Cat#H4034** |
| ifosfamide | **Shionogi** | |
| isoflurane | **Pfizer** | |
| KSOM medium | **ARK Resource** | |
| levofloxacin | **Daiichi-Sankyo** | |

| | | |
|---|---|---|
| Metacam | **Boehringer Ingelheim** | |
| metronidazol | **Pfizer** | |
| mWM medium | **ARK Resource** | |
| nifekalant | **Astellas** | |
| Opti-MEM | **Thermo Fisher** | **Cat#11058021** |
| PrimeScript RT | **Takara** | **Cat#RR047A** |
| Procainamide | **alfresa** | |
| RNeasy | **Qiagen** | **Cat#74106** |
| TaqMan Fast Advanced Master Mix | **Thermo Fisher** | **Cat#4444557** |
| Tissue ATP Assay Kit | **Toyo INK** | **Cat#TA100** |
| vanoxerine | **MCE** | **HY-13217A** |
| verapamil | **Eisai** | |
| Williams' medium E | **Thermo Fisher** | **Cat#A1217601** |

| | | |
|---|---|---|
| **Software and Algorithms** | | |
| Image J | **NIH** | **https://imagej.nih.gov/ij/** |
| MetaMorph | **Molecular Devices** | |
| StackReg (Image J plugin) | **EPFL** | **http://bigwww.epfl.ch/thevenaz/stackreg/** |
| **Oligonucleotides** | **eurofins** | |
| Sequences written 5´–3´ (one sequence/line)<br>P1 AGAGCCTCTGCTAACCATGTTCATGCCTTC<br>P2 GTGACACTAAGTCAAACGCGAAA<br><br>for Kusabira Orange<br>5’-AGAGATGACACTACGCGTCACAA,<br>5’- GTGACACTAAGTCAAACGCGAAA,<br>the TaqMan probe 5’- CCGAGGGCGGGCCAATGC.<br><br>for the internal control sequence, *Gnrhr*<br>5’-TGTATGCCCCAGCTTTCATG<br>5’-GGCTGAGTGATGGCCAGG<br>the TaqMan probe 5’-TGGTGGTGATTAGCCTGGACCGCT | | |

### CONTACT FOR REAGENT AND RESOURCE SHARING

Further information and requests for resources and reagents should be directed to and will be fulfilled by the Lead Contact, Masamichi Yamamoto

### EXPERIMENTAL MODELS AND SUBJECT DETAILS

#### Imaging of Muscle Contraction and Measurement of Torque

Mice were anesthetized (isoflurane, 1.5%–2.0% using a ventilator), the leg hair was shaved, and skin was stripped from the leg. To detect fluorescence, the leg was immobilized in a custom restraint and kept warm using a stage heater (Tokai Hit). The exposed tibialis anterior (TA) muscle was observed using a fluorescence stereo microscope. To measure torque, each mouse was positioned with the right foot on the footplate with the ankle joint angle positioned at 90°. During muscle contractions, isometric dorsi-flexional torque was measured, and to avoid the influence of plantar flexion, the Achilles tendon was severed. Muscle contractions were induced by electrical stimulation of the sciatic nerve (10, 20, 30, 40, 60, 100 Hz).

#### GO-ATeam Mouse Model of Myocardial Infarction

Mice were anesthetized (isoflurane, 1.5%–2.0% using a ventilator), intubated, and ventilated. The left anterior descending artery was accessed by lateral thoracotomy and pericardectomy. Left-anterior descending artery ligation was performed using an 8-0 suture placed caudal to the left atrial appendage. Ligation was confirmed by blanching of the myocardium and electrocardiogram (ECG). The wound was closed in layers using a 6-0 suture and Aron Alpha A. Mice were closely monitored after surgery, and meloxicam (Metacam) or buprenorphine was administered as required for analgesia.

**Drug-Induced Cardiotoxicity in GO-ATeam Mice**

Mice were anesthetized (isoflurane, 1.5%–2.0% using a ventilator), intubated, and ventilated. Drug administration was performed by placing a 30G needle connected to a syringe pump (15 μl /minutes) in the right jugular vein while performing an ECG for 60 minutes. The concentration of each drug was the following: furosemide (10mg/ml); doxorubicin (2mg/ml); 5-FU (50mg/ml); ifosfamide (10mg/ml); disopyramide (0.1mg/ml); nifekalant (1.25mg/ml); verapamil (0.25mg/ml); levofloxacin (25mg/ml); alfuzosin (2mg/ml); vanoxerine (0.25mg/ml); cyclophosphamide (20mg/ml); procainamide (0.25mg/ml); droperidol (0.125mg/ml); erythromycin (0.5mg/ml); amphotericin B (4mg/ml); azithromycin (5mg/ml); ciprofloxacin (0.4mg/ml); metronidazol (1mg/ml). With doxorubicin, the intrinsic fluorescence is emitted as red light, so the difference value was used as the background. Therefore, the ratio image of doxorubicin does not show the correct amount and was omitted from Fig. S6A.

### METHOD DETAILS

#### Cell Lines

Mouse embryo fibroblasts were generated from embryonic 15.5 day embryos (Nagy, 2003). Mouse embryos (15.5-days postcoitus) were dissected in a petri dish containing PBS, and the limbs, internal organs, and brain were removed; the embryos were then rinsed three times with DMEM without serum. The embryos were minced into very small pieces using sterile surgical scissors. The minced embryos were added to a tube containing trypsin/EDTA in PBS, incubated at 37ºC for 30 min, and centrifuged at 200x*g* for 5 min. The cells were then added to a tissue culture dish containing DMEM plus 10% FBS.

#### Tissue Culture

Livers were cultured in a flow chamber containing Williams' medium E without phenol-red. Ibuki #180 air stones (Ibuki, Japan) were used to bubble in 95% $O_2$ and 5% $CO_2$.

#### Embryo culture

One-and two-cell embryos were cultured in KSOM medium without phenol red in 5% $CO_2$.

#### Live Imaging of Embryos

We used warm KSOM medium without phenol red to flush two-cell stage embryos from the uteri of pregnant knockin mice. Embryos were placed in no. 1 glass-bottom dishes (Matsunami Glass Ind., Ltd.) filled with KSOM without phenol red, which were placed in a humidified cell culture incubator (Tokai Hit) set to 37°C and provided with a continuous supply of 5% $CO_2$. Embryos were observed using an inverted multiphoton microscope (TCS SP8 MP; Leica) equipped with an HC FLUOTARL 25×/NA0.95 water objective (Leica). The embryos were treated with 10 mM 2-deoxy-D-glucose (Wako) to inhibit glycolysis and 36 μM antimycin A (Sigma-Aldrich) to inhibit oxidative phosphorylation (OXPHOS).

**Animal model**

Animals were housed (5 animals at maximum in each cage) and maintained under a 12-h light: 12-h dark cycle, at constant temperature and humidity (20-24℃, 35-55%), with food and water *ad libitum*. All animal experiments were performed in accordance with protocols approved by the Committee of Experimental Animal Research of Gunma University, Kyoto University, and Nagoya Institute of Technology, and were approved by the Ethical Committee of the Gunma University, Kyoto University, and Nagoya Institute of Technology. All animal model and experimental procedures conformed to the NIH Guide for the Care and Use of Laboratory Animals.

**Intravital Imaging of Organs**

Mice were anesthetized (ventilation using 1.5%–2.0% isoflurane; Narcobit-E and KN-58 SLA; Natsume Seisakusyo), intubated, ventilated, and continuously monitored using ECG. The body hair was shaved using an electric clipper. The thoracic and peritoneal cavities were exposed using electrical cautery, and the mouse was immobilized in a custom restraint and kept warm using a stage heater (Tokai Hit). Meloxicam (Metacam) or buprenorphine was administered as required for analgesia.

A catheter was inserted into the jugular vein using a 30-gauge needle attached to PE-10 tubing (Becton Dickinson). Organs were observed using an inverted multiphoton microscope (TCS SP8 MP; Leica) equipped with an HC FLUOTARL 25×/NA0.95 water objective (Leica) or a fluorescence stereo microscope (M165FC; Leica) equipped with a PLAN APO 1.0× objective (Leica).

**MALDI-Imaging Mass Spectrometry**

Liver tissues were dissected and embedded in Super Cryoembedding Medium (SCEM)-L1 (SECTION LAB, Hiroshima, Japan), and stored at –80°C until use. The frozen SCEM blocks were sectioned at –16°C using a cryostat (CM 3050; Leica, Wetzlar, Germany) to make 8-µm thick liver slices that were thaw-mounted onto indium-tin-oxide (ITO)–coated glass slides (Bruker Daltonics, Billerica, MA, USA). Tissue sections were coated with 2,5-dihydroxybenzoic acid (50 mg/mL dissolved in 80% methanol) (Benabdellah et al., 2009; George et al., 2007) as a matrix for cationic molecular detection. This solution was manually sprayed with an artistic-brush (Procon Boy FWA Platinum, Mr. Hobby, Tokyo, Japan). MALDI imaging was performed using an Ultraflextreme MALDI-TOF/TOF mass spectrometer (Bruker Daltonics) for anion metabolite imaging. For TOF/TOF-MS-based imaging analysis, data were acquired in the negative reflectron mode, and signals between *m/z* 50 and 1000 were collected.

Analysis and image reconstruction was performed using FlexImaging 4.1 software (Bruker Daltonics).

**Sample preparation for metabolome analysis**

Metabolite extraction from tissues for metabolome analyses was performed as described previously(Miyazawa et al., 2017; Oka et al., 2017). Briefly, liver tissues, together with internal standard (IS) compounds (see below), were homogenized in ice-cold methanol (500 μL) using a manual homogenizer (Finger Masher (AM79330), Sarstedt), followed by the addition of an equal volume of chloroform and 0.4x volume of ultrapure water (LC/MS grade, Wako). The suspension was then centrifuged at 15,000x*g* for 15 min at 4°C. After centrifugation, the aqueous phase was ultrafiltered using an ultrafiltration tube (Ultrafree MC-PLHCC, Human Metabolome Technologies). The filtrate was concentrated using a vacuum concentrator (SpeedVac, Thermo). The concentrated filtrate was dissolved in 50 μL of ultrapure water and used for LC-MS/MS and IC-MS analyses.

**Quantification of metabolites using internal and external standards**

We used an internal standard (added to the tissue before extraction) and an external standard to determine *m/z* values and specific retention times of LC or IC for all metabolites. The detailed method is as follows: internal standards were 2-morpholinoethanesulfonic acid (MES) and 1,3,5-benzene-tricarboxylic acid (trimesate) for anionic metabolite measurements and L-methionine sulfone and 3-aminopyrrolidine dihydrochloride were used for cationic metabolite analysis. These compounds are not present in the tissues. Thus, they serve as ideal standards. Loss of endogenous metabolites during sample preparation was corrected by calculating the recovery rate (%) for each sample measurement. Before sample analysis, we measured the mixture of authentic target metabolites (external standards) in ultrapure water to determine *m/z* values and retention times of all metabolites.

For metabolome analysis of anionic metabolites of glycolysis, the TCA-cycle, and the PPP as well as nucleotides, we used an orbitrap-type MS (Q-Exactive focus, Thermo Fisher Scientific), connected to a high performance ion-chromatography system (ICS-5000+, Thermo Fisher Scientific). This system was used to perform highly selective and sensitive metabolite quantification afforded by IC-separation and the Fourier Transfer MS principle (Hu et al., 2015). The IC was equipped with an anion electrolytic suppressor (Thermo Scientific Dionex AERS 500) to convert the potassium hydroxide gradient into pure water before the sample entered the mass spectrometer. The separation was performed using a Thermo Scientific Dionex IonPac AS11-HC, 4-μm particle column. The IC flow rate was 0.25 mL/min supplemented postcolumn with 0.19 ml/min makeup flow of MeOH. The potassium hydroxide gradients for IC separation were as follows: from 1 mM to 100 mM (0–40 min), 100 mM (40–50 min), and 1 mM (50.1–60 min) (column temperature, 30°C).

The Q Exactive focus mass spectrometer was exclusively operated under ESI-negative mode. A full mass scan (*m/z* 70−900) was performed at a resolution of 70,000. The automatic gain control (AGC) target was set to $3 \times 10^6$ ions, and the maximum ion injection time (IT) was 100 ms. Source ionization parameters were optimized using a 3-kV spray voltage. Other relevant parameters were as follows: transfer temperature, 320°C; S-Lens level, 50; heater temperature, 300°C; sheath gas, 36°C; and aux gas, 10°C.

The amount of cationic metabolites in liver tissues was quantified using liquid chromatography-tandem mass spectrometry (LC-MS/MS). Briefly, a triple-quadrupole mass spectrometer equipped with an electrospray ionization (ESI) ion source (LCMS-8040, Shimadzu Corporation) was used for positive- and negative-ESI and multiple reaction monitoring (MRM) modes. The samples were resolved using the Discovery HS F5-3 column (2.1-mm i.d. x 150 mm, 3-μm particle; Sigma-Aldrich), using a step gradient with mobile phase A (0.1% formate) and mobile phase B (0.1% acetonitrile) at ratios of 100:1 (0–5 min), 75:25 (5–11 min), 65:35 (11–15 min), 5:95 (15–20 min), and 100:0 (20–25 min) at 0.25 ml/min (column, 40°C).

**Organ slices**

Mice were anesthetized using isoflurane 4.0%. These livers were soaked in cooled Williams' medium E without phenol-red, and Ibuki #180 air stones (Ibuki, Japan) were used to bubble in 95% $O_2$ and 5% $CO_2$ or 95% $N_2$ and 5% $CO_2$ to induce hypoxia. A vibratome slicer (VT 1000S; Leica) was used to prepare 300-μm-thick liver slices that were immediately placed into the chamber, supplied with Williams' medium E, which was placed on the stage of the microscope. Liver slices were observed using an inverted multiphoton microscope or fluorescence stereo microscope.

**Imaging Conditions**

The imaging system included a TCS SP8 MP multiphoton microscope (Leica), driven by a Mai Tai HP Ti:Sapphire laser (Spectra-Physics) tuned to 920 nm and an inverted microscope equipped with an HC FLUOTARL 25×/NA0.95 water objective (Leica). Cells expressing GO-ATeam were detected using a bandpass emission filter, 525/50 nm for EGFP and 585/40 nm for Kusabira Orange. A second imaging system included an M165FC fluorescence stereo microscope equipped with a PLAN APO 1.0× objective (Leica). Organs were exposed to excitation light (ET470/40), and images were captured using a dual-view fluorescence cMOS Camera (ORCA-Flash 4.0, Hamamatsu photonics, Japan). We used D515/30, DM540, and D575/40 filters to resolve fluorescence emission peaks.

All experimental procedures performed according to the guidelines for the Care and Use of Animals for Experimental Procedures of Gunma University and Kyoto University and were approved by the local committees of the Graduate Medical School, Gunma University and Kyoto University that regulate the handling of experimental animals.

**Image Processing**

Images of Kusabira Orange EGFP fluorescence were analyzed using MetaMorph software (Molecular Devices). ImageJ was used for video editing and linear adjustment of intensity, only for visualization. To eliminate the XY drift in time-lapse series, we used the StackReg plugin for ImageJ. To determine the fluorescence intensities of EGFP and Kusabira Orange at 16 bits, we used MetaMorph and ImageJ.

**Measurement of ATP Levels in Mouse Embryonic Fibroblasts (MEFs)**

MEFs (Nagy, 2003) derived from the knockin mouse embryos were permeabilized by exposing them to 50 mg/mL alpha-hemolysin (Sigma-Aldrich) dissolved in Permeabilization Buffer (140 mM KCl, 6 mM NaCl, 0.1 mM EGTA, and 10 mM HEPES (pH 7.4)) for 30 min in a $CO_2$ incubator. Next, permeabilization buffer was replaced with calibration buffer (140 mM KCl, 6 mM NaCl, 0.5 mM $MgCl_2$, and 10 mM HEPES, pH 7.4) that contained different concentrations of Mg-ATP (0–20 mM). The permeabilized MEFs were observed using an inverted multiphoton microscope.

**Luciferase Assay**

Luciferin-luciferase detection of ATP was performed using a Tissue ATP Assay Kit (c B-NET) according to the manufacturer's protocol. Briefly, weighed, fresh tissue samples were homogenized in cold homogenization buffer (0.25 M sucrose and 10 mM HEPES, with the pH adjusted to 7.4 by NaOH). One sample was distributed into three tubes, and luciferase reagent (L1) was added immediately before measurements using a luminometer (Lumate, LB-9507, Berthold Technologies). A standard curve was generated daily before each measurement using standards and ATP-free water provided in the kit. The concentrations are expressed per milligram of wet tissue.

**Electroporation**

A pair of custom (BEX, Tokyo, Japan) platinum-block electrodes (length, 10 mm; width, 3 mm; height, 0.5 mm; gap, 1 mm) was used. The electrodes, which were connected to a CUY21EDIT II unit (BEX, Tokyo, Japan), were placed under a stereo microscope. Embryos cultured in mWM medium (ARK Resource, Kumamoto, Japan) were washed three times with Opti-MEM (Thermo Fisher Scientific) to remove the serum-containing medium. The embryos were then placed in a line in the electrode gap filled with RNA-containing Opti-MEM I solution (total 5 μl), and electroporation was then performed. The electroporation conditions were 30 V (3 ms ON + 97 ms OFF) × 7 in most experiments. After electroporation, the embryos were immediately collected from the electrode chamber and subjected to four washes with M2 medium (Sigma-Aldrich) followed by two washes with mWM medium. The eggs were then cultured in mWM medium at 37 °C in an atmosphere containing 5% $CO_2$.

### QUANTIFICATION AND STATISTICAL ANALYSIS

Data are expressed as the mean ± SEM. Statistical analysis was performed using the unpaired two-tailed Student *t* test to compare two groups and analysis of variance (ANOVA) (*$P < 0.05$; **$P < 0.01$; NS, not significant). All data were normally distributed, and variance was similar between groups. Biological replicates were performed using different samples derived from different mice. The results represent data from at least three independent experiments. We estimated the required sample size considering the variation and mean of the samples. We used the fewest animals required for statistically valid conclusions. Our protocol required excluding mice if we observed abnormal size, weight, or both or disease symptoms before performing experiments. However, this was unnecessary, as all mice were phenotypically normal and healthy.

### SUPPLEMENTAL FIGURES & FIGURE LEGENDS

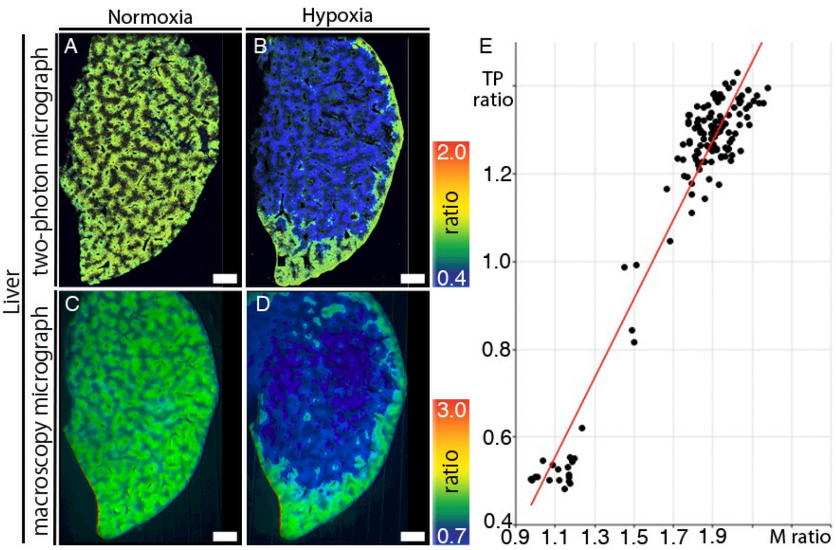

Figure S1

**Fig. S1. Strong Positive Correlation of FRET/GFP Ratios Between Two-Photon Images and Low Magnified Images, Related to Fig. 1**

Liver slices (A–E) of GO-ATeam mice. (B, D) Images of the tissues shown in A and C after the slices were exposed to hypoxic conditions. FRET/GFP ratios from 0.4 to 2.0 in two-photon images (A, B) and from 0.7 to 3.0 in low magnified images (C, D). (E) FRET/GFP ratio correlation diagram. The ratios of two-photon microscope images on the x-axis are plotted as a function of the ratios of low magnified images on the y-axis. Correlation coefficients ($R^2 = 0.92$, n = 139). Scale bars indicate 1mm (A-D).

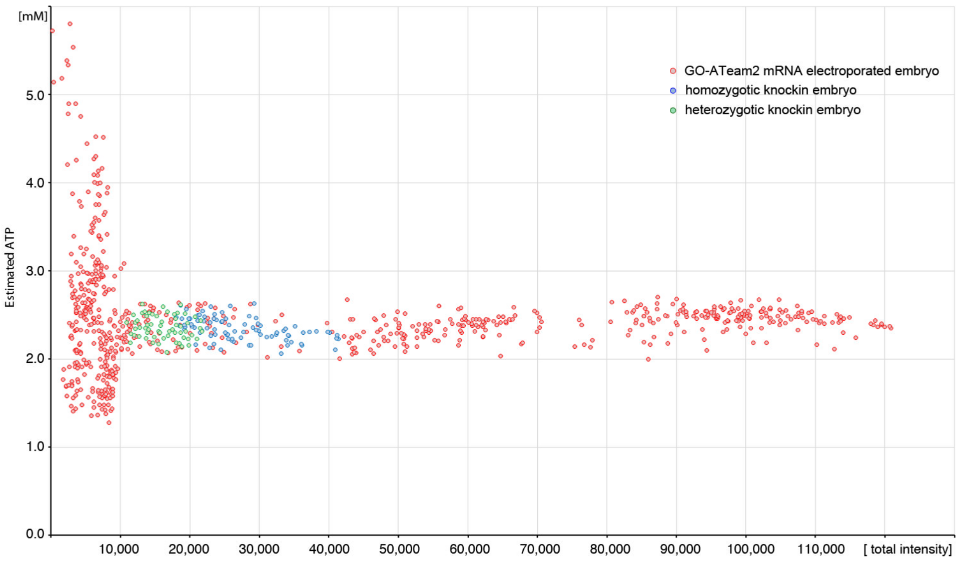

Figure S2

**Fig. S2. Correlation between the FRET/GFP Ratio and Total Intensity in Two-Cell Embryos, Related to Figs. 1, 2**

Embryos were collected from pregnant wild-type and GO-ATeam heterozygous (green dots, n = 88) or homozygous mice (blue dots, n = 68). Wild-type embryos were electroporated with GO-ATeam2 mRNA, cultured overnight, and observed using a two-photon microscope (red dots, n = 756). ATP concentrations (mM) calculated from the FRET/GFP ratios (y-axis) are shown as a function of the sum of the average fluorescence intensities of the GFP plus FRET signals that represent total average fluorescence intensities acquired at 16 bits on the horizontal axis.

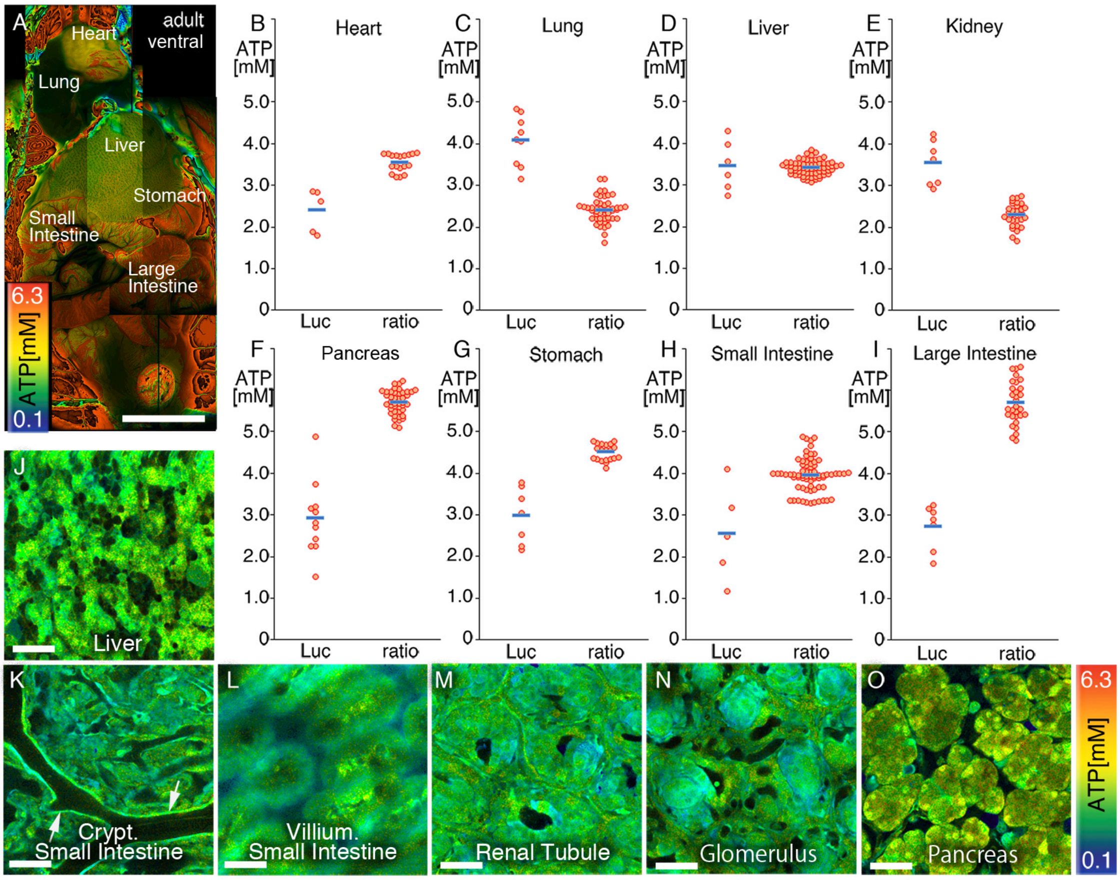

Figure S3

**Fig. S3. ATP levels in GO-ATeam Live Mice and Intravital Organs, Related to Fig. 2**

(A) FRET/GFP fluorescence of adult (aged 8 weeks) GO-ATeam mice.

(B–I) ATP concentrations (luciferase assay, “Luc”) and the FRET/GFP ratios (right, “ratio”) in heart (B), lung (C), liver (D), kidney (E), pancreas (F), stomach (G), small intestine (H), and large intestine (I) in adult GO-ATeam mice.

Intravital FRET/GFP imaging of postnatal day-0 [J–O] GO-ATeam mice. Liver (J), small intestine (K, L, arrow: blood vessel), kidney (M, N), pancreas (O). Scale bars indicate 10mm (A), or 100µm (J-O).

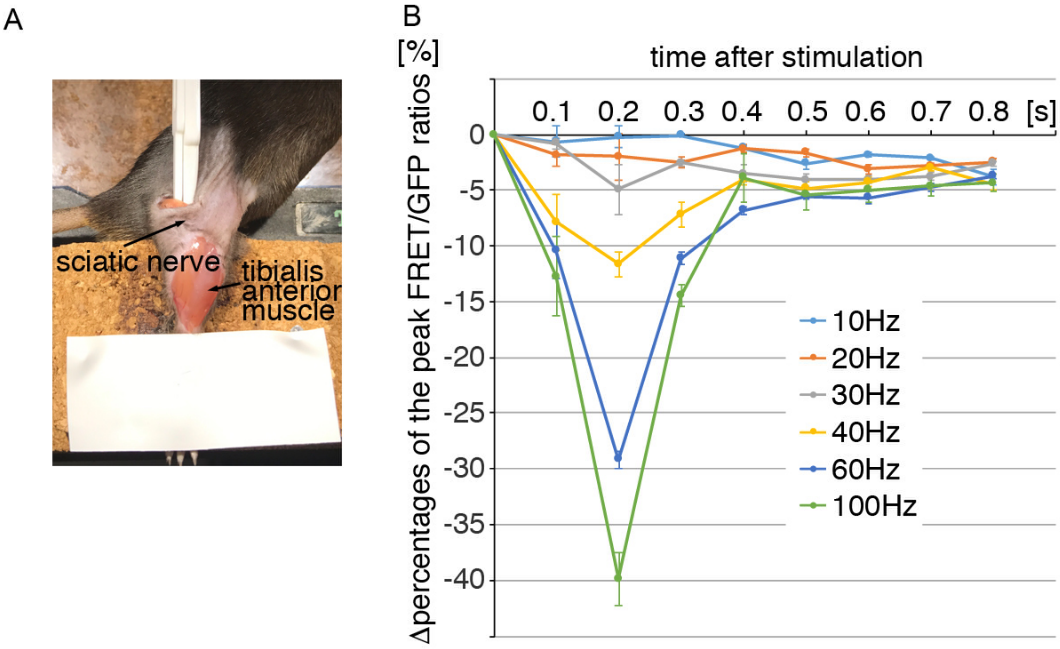

Figure S4

**Fig. S4. ATP Dynamics Correspond to the Force Generated by Muscles, Related to Fig. 3**

(A) Method for observing the tibialis anterior muscle by dissecting the sciatic nerve. (B) The delta percentages of the peak FRET/GFP ratios (n = 4) as a function of time after stimulating the sciatic nerve.

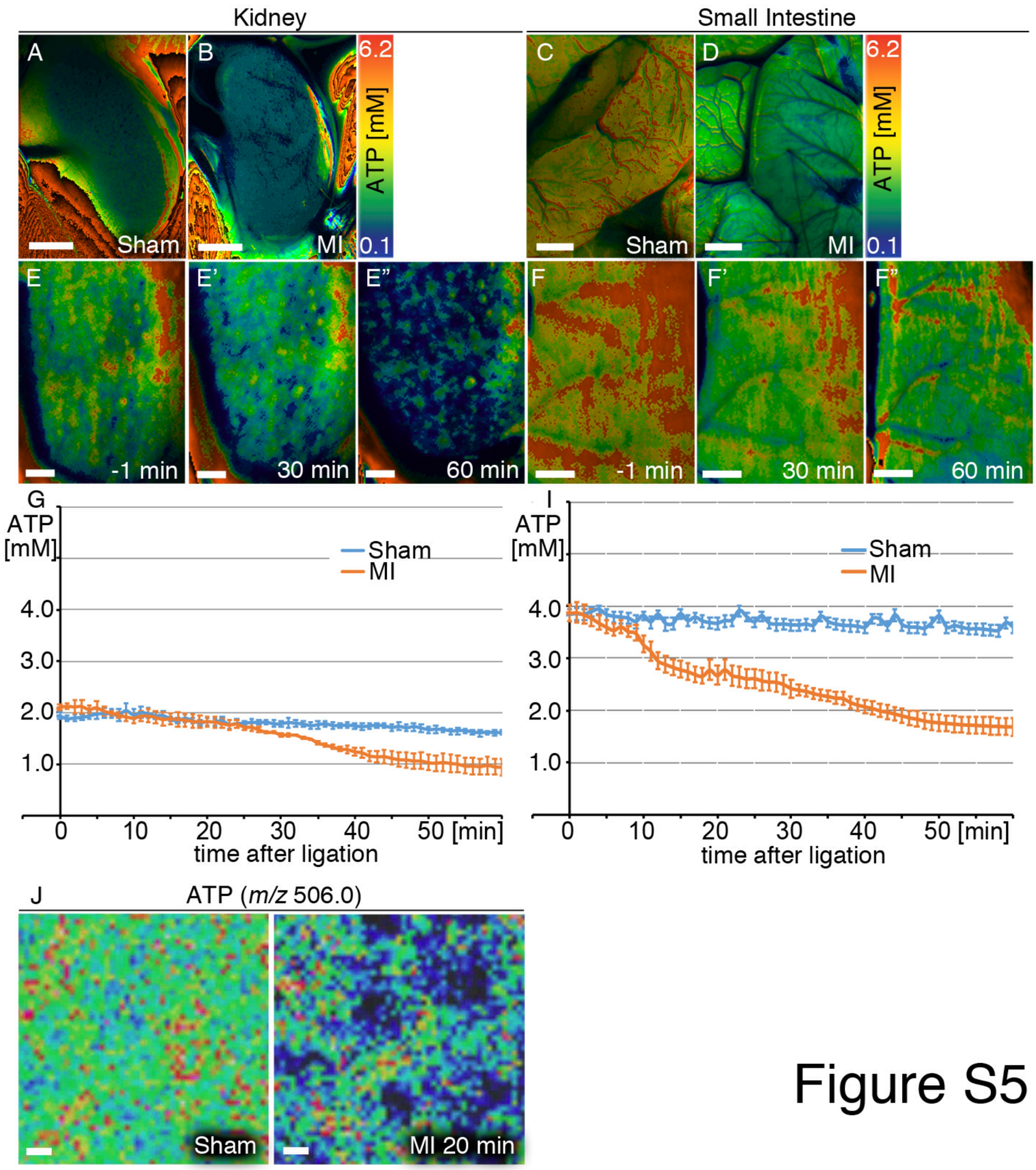

**Fig. S5.**

**Effects of Myocardial Infarction on the ATP Concentrations of GO-ATeam Mice, Related to Fig. 4**

Intravital images of FRET/GFP ratios 5 days after ligation of the left anterior descending artery (LAD) (myocardial infarction, MI) (B, D), or control sham-operated mice (sham) (A, C). Intravital time-lapse images and graphs of FRET/GFP ratios in organs (E–I) after ligation of the LAD. Kidney (A, B, $n = 8$ each; E–E”, G, $n = 4$ each, after $t = 32$ min, $p<0.05$) and small intestine (C, D, $n = 8$ each; F-F”, I, $n = 6$ each, after $t = 12$ min, $p<0.05$).

(J) Imaging Mass Spectrometry of ATP (*m/z* 506.0) in the liver sham-operated mice, and 20 min after ligation of the LAD. Scale bars indicate 2mm (A-D), 1mm (E-F”), or 100μm (J).

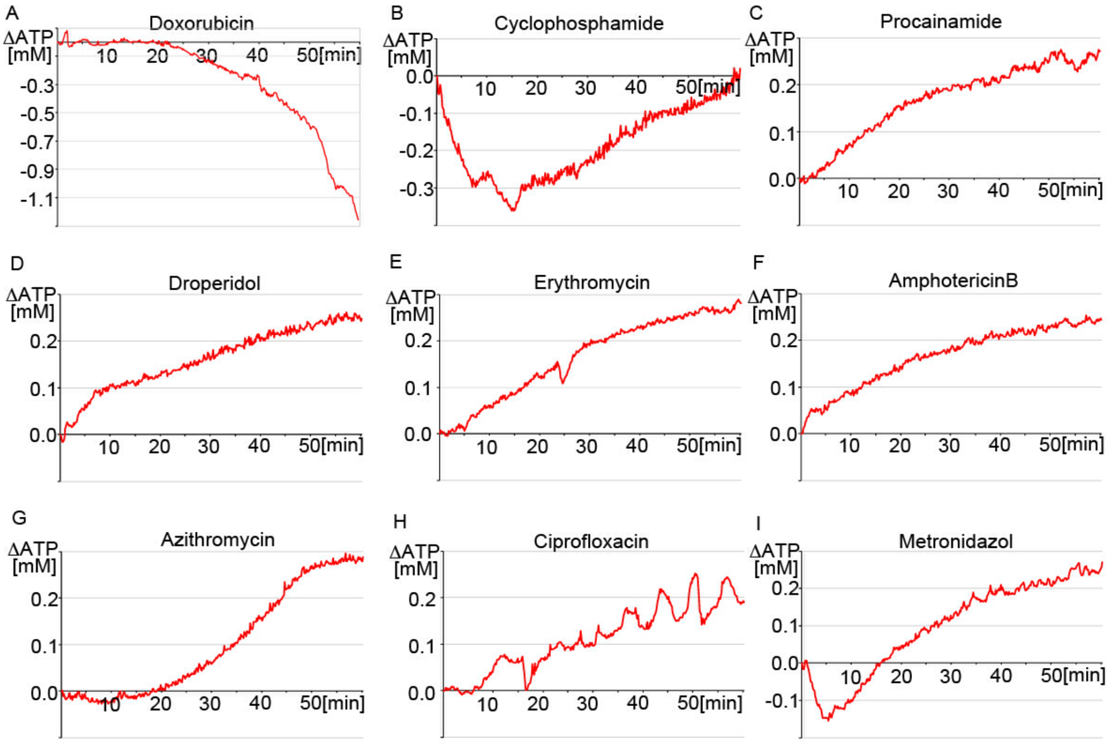

Figure S6

**Fig. S6. Effects of Drug-Induced Cardiotoxicity on ATP Concentration in the Heart, Related to Fig. 5.**

(A-I) Intravital time-lapse imaging of ATP concentrations calculated from the FRET/GFP ratios in the heart using the fluorescence stereo microscope. The graph shows the change ATP level (y-axis) after administration (red line, indicated drug). Horizontal axis shows time [minutes] after administration. (A) doxorubicin (n=3) (B) cyclophosphamide (n=4), (C) procainamide (n=5), (D) droperidol (n=9), (E) erythromycin (n=7), (F) amphotericin B (n=4), (G) azithromycin (n=5), (H) ciprofloxacin (n=5), (I) metronidazol (n=3).
